# Volumetric assessment and longitudinal changes of brain structures in formalinized Beagle brains

**DOI:** 10.1101/2021.12.03.471167

**Authors:** Francesca Del Signore, Germain Arribarat, Leonardo Della Salda, Giovanni Mogicato, Alexandra Deviers, Benjamin Cartiaux, Massimo Vignoli, Patrice Peran, Francesco de Pasquale

## Abstract

High field MRI represents an advanced technique both for diagnostic and research purposes on animal models such as the Beagle dog. The increasing interest in non-invasive neuroscience, aging, and neuropathological research led to a need of reference values (in terms of volumetric assessment) for the typical brain structures involved and, nowadays, several canine brain MRI atlases have been provided. Since no reports are available regarding the measurements reproducibility and few information are available about formalin fixation effect on brain structures when applied to MRI segmentation, we assessed the segmentation variability of selected structures as a function of the operator (two operators segmented the same data) and their intrinsic variability within a sample of 11 Beagle dogs (9 females and 2 males, 1.6 ± 0.2 years). Then, we analyzed for one further Beagle dog (2 years old) the longitudinal changes in the brain segmentations of these structures corresponding four conditions: *in vivo, post mortem* (after euthanasia), *ex vivo* (brain extracted and studied after 1 month in formalin and after 11 months); all the MRI images were collected with a 3 T MRI scanner. Our findings suggest that the segmentation procedure can be considered overall reproducible since only slight statistical differences were detected, apart from the ventricles.

Furthermore, in the *post mortem/ ex vivo* comparison, the majority of the structures showed a higher contrast leading to more reproducible segmentations across operators and a net increase of volume of the studied structures; this could be justified by the intrinsic relaxation time changes observed as a consequence of formalin fixation, that led to an improvement of brain structures visualization and then segmentation.

To conclude, MRI based segmentation seems to be a useful and accurate tool that allows longitudinal studies, especially when applied to formalin fixed brains.

## Introduction

High field MRI, thanks to its high SNR and short acquisition time, represents an advanced technique both for diagnostic and research purposes on animal models such as the dog [1,2]. This model offers several advantages over more standard rodent and primate ones as testified by the growing literature on neurocognitive, aging, and clinical applications [3]. Neurocognitively, the canine shares similar behavioral/emotional responses with humans, e.g. in linking learning, memory, and other cognitive functions. These convergent sociocognitive skills places the dog in a unique position to increase our understanding of sociocognitive in humans [4]. Being gyrencephalic, the dog brain, as compared to rodent and avian, it represents a better experimental model for several disorders, e.g. gliomas and aging [5–7]. Among domestic canines, the Beagle is the breed most commonly used in laboratories, thanks to its moderate size, docile nature, and strong immunity [8–10]. Such increasing interest in non-invasive neuroscience, aging, and neuropathological research led to a need of reference values (in terms of volumetric assessment) for the typical brain structures involved. To this aim, recent studies developed a standard canine brain atlas to provide a common spatial referencing and architectonic-based cortical segmentation of coregistered data [4]. Similarly, a stereotactic cortical atlas for the mesaticephalic canine brain has been recently developed for functional and structural MRI analyses [3]. In general, the currently available atlases, see for example [11–13], are affected by some limitations such as the small number of subjects used [11], the acquisition of non-isovolumetric data [13], the use of dogs non neurologically/clinically healthy [12] and the mixing of different breeds included in the sample [3]. To overcome the uncertainty related to the breed variability, very recently Liu et al. realized a specific atlas for the Beagle breed, which is the most used breed in this kind of studies [14]. However, all these studies were performed on alive subjects, apart from the work of Datta and colleagues, 2012, where formalin-fixed brains (*ex vivo*) were segmented to obtain a diffeomorphic brain atlas of mesaticephalic dogs coregistered onto an *in vivo* template [11]. Nevertheless, in this study, some important aspects such as the reproducibility of the measurements across different operators and the volumetric variation from *in vivo* to *ex vivo* phases were not assessed [11]. Such a relationship is very important since MRI findings can be linked and often validated through histopathology. This can be very time-consuming and challenging for several reasons, e.g. the inaccurate correspondence MRI-anatomical sections (due to different slice thickness and orientations) [15]. For this reason, when whole-brain histopathology is not feasible, *in vivo* and *post mortem* MRI can be used as a guide for limited pathological sampling [16–18]. Similarly, in forensic radiology, *post mortem* MRI has been recognized as a supplementary tool to address specific forensic questions [19–21]. However, *ex-vivo* MRI can be very challenging. First, after death, the brain undergoes several changes, e.g. microbial degradation, autolysis, breakdown of cell membranes, and stochastic diffusion of molecules. Second, also the chemical fixation, needed to ensure the longitudinal stability of the macromolecular structure, might affect the tissue properties. Therefore, due to death and fixation, a series of artifacts and changes of tissue properties are expected. This will affect MR imaging and the conclusions based on MRI measurements in fixed tissue may not reflect directly the *in-vivo* environment [21]. For example, it has previously been reported that formalin fixation causes a tissue shrinkage that not be homogeneous among the various brain structures, and it might vary over time [22]. However, these aspects are still under debate and the literature is scares. For this reason, in this study, we analyzed the effect of long-term fixation (12 months) on brain structures in a sample of 11 Beagle dogs. We manually segmented a set of brain structures, e.g. Globus Pallidus, Caudate, and Substantia Nigra, whose volumetric changes have been reported to correlate with many neurodegenerative disorders such as Parkinson’s [23]. First, we assessed the variability of the extracted volumes as a function of the operator (two operators segmented the same data) and their intrinsic variability within the sample. Then, we analyzed for one further dog the longitudinal changes in the brain segmentations of these structures corresponding four conditions: *in vivo, post mortem* (after euthanasia), *ex vivo* (brain extracted and studied after 1 month in formalin and after 11 months). In this way, the last condition overlaps with the previous sample of 11 dogs.

The estimated volumetric data can represent an important reference for future studies and the longitudinal effects observed might shed light on which structures are segmented more accurately as a function of time spent in formalin. As far as we know, this is the first study reporting brain structures in formalinized dogs and their longitudinal changes.

## Materials and Methods

### Animals

A sample of 12 healthy Beagle dogs were evaluated in two studies. In the first study, a group of 11 dogs (9 females and 2 males, 1.6 ± 0.2 years) was used for the evaluation of the effect of long-term fixation on MRI properties of the brain. In this context, a single MRI scan was performed on isolated heads that remained fixed for 11 months. Dogs originated from a laboratory in which they completed their research time and were euthanized for teaching purposes (i.e. preparation of veterinary anatomical teaching materials: MRI brain atlas and embalmed cadavers for dissection).

In the second study, one dog (male, 2 years) was used for the longitudinal evaluation of the effect of death and fixation on MRI properties of the brain. This dog underwent 4 MRI exams: 1 *in vivo*, 1 *post-mortem* performed just after euthanasia, and 2 *ex vivo* performed on the brain removed from the skull after 1 (*ex_vivo_1*) and 12 months of fixation (*ex_vivo_12*). While the terms *post-mortem* and *ex vivo* are normally interchangeable, in this study, they refer to two different conditions which are evaluation of the brain confined by the skull just after death (*post-mortem*) and evaluation of formalin-fixed brains (*ex vivo*, either isolated or confined by the skull).

The experimental procedures related to the preparation of veterinary anatomical teaching materials were approved by the Animal Ethics Committee of the National Veterinary School of Toulouse with authorization n° 21559-2019071917392588. Dogs were euthanized by an IV injection of ≥100 mg/kg sodium pentobarbital while they were deeply anesthetized (anesthetic protocol: IV injection of butorphanol (0,4 mg/kg), medetomidine (20 μg/kg), and diazepam (0,2 mg/kg)). Heparin sodium (1000 IU) was injected by IV route 5 minutes before euthanasia to optimize post-mortem perfusion of fixative solution.

### Fixation protocol

Dogs were anesthetized to acquire in vivo MR images (not used in the present study) after which they were euthanized, and their heads were fixed according to the procedure described above. Heads were then stored in containers filled with 10% formalin solution and were scanned after 11 months of fixation.

For the dog in the second study, in vivo MRI scans and euthanasia were carried out under anesthesia with the protocol described above. The post-mortem MRI examination was performed just after euthanasia and once this acquisition was completed, the cadaver was transferred to a special room for fixation. The head was then separated from the body to be perfused via the common carotid arteries with a rinsing solution (NaCl, flow rate: 15 mL/minute, perfusion time: 5 minutes) and a fixative solution (10% formalin solution, 15 mL/minute, perfusion time: 5 minutes). The head was stored in a container filled with 10% formalin solution. After one month of fixation, the brain was removed from the skull for an ex vivo MRI acquisition. An additional ex vivo MRI examination of the brain was performed after 11 months of fixation.

### MRI acquisition

MRI examinations were performed at the Institute for Brain Sciences of Toulouse using a high field 3.0 Tesla magnet (Philips ACHIEVA dStream) at the Inserm/UPS UMR1214 ToNIC Technical Platform, and a, 8-channel, human elbow coil (serving as dog head coil) for signal reception. The Ex-vivo examinations were performed with a 1-channel solenoid antenna. To guarantee a homogeneous and accurate signal, no acceleration or preparation factors were used. For the dog with longitudinal follow-up of the brain, the imaging protocol comprised of T1and T2 weighted images. The 3D whole-brain T1 and T2 weighted images were obtained in the sagittal plane. For T1 imaging (Fast Field Echo), the sequence parameters were the following: echo time/repetition time =4.0/9.0 ms, flip angle = 8°. For T2 imaging (Spin Echo) the sequence parameters were the following: echo time/repetition time = 266/2500 ms, flip angle = 90°. The spatial resolution parameters were the same for both acquisitions: pixel spacing 0.5 × 0.5 mm^2^, slice thickness = 0.5 mm, matrix size = 288×288, numbers of slices = 300, voxel size = 0.5 x 0.5 x 0.5 mm3. The total duration of the imaging protocol was 60 min.

These sequences were used for the 4 MRI examinations of the follow-up. Twenty-four hours before the ex vivo MRI scans, the brain was rinsed with water and then submerged in a 0,9% saline solution (NaCl). Just before acquisition, the brain was put in an MRI-compatible container (a plastic container with a leakproof screw cap) totally filled with saline solution, the container was then placed in the elbow coil.

Twenty-four hours before their acquisition, the 11 fixed heads were rinsed with water and then submerged in NaCl. For the acquisition, they were dried, wrapped up in hermetic packages, held horizontally on the MRI table, and placed in the human elbow coil. The imaging protocol comprised T1-weighted images (repetition time TR = 8.5 ms; echo time TE = 3.8 ms; voxel size 0.5×0.5×0.5 mm, matrix 288×288×300) and T2-weighted images (repetition time TR = 265.71 ms; echo time TE = 2500 ms; voxel size 0.5×0.5×0.5 mm, matrix 288×288×300).

### Assessment of inter-operator reproducibility

Two veterinarians experienced in canine brain structure segmentation operators (O1- and O2 respectively) blindly segmented the following structures: Ventricles (from T1 scans) and Caudate Nucleus, Hippocampus, Substantia Nigra, Putamen, Globus Pallidus, Lateral and Medial Geniculate Nucleus (from T2 scans), dividing the measurements in left and right parts on the group of 11 dogs and the longitudinal study. The choice to perform the segmentations either in T1 or T2 was driven by the contrast observed in the various structures.

As far as it regards the analysis on the measurements collected on the group of 11 dogs, on each structure mean (M) and standard deviation (SD) was evaluated. Based on them, to assess the intra-operator reproducibility, the percentage volumes were computed as CV = SD/M. Finally, the statistical comparison between the two operators was provided by a t-test.

### Longitudinal study

For the second part of the study, the two operators performed the same segmentations of the previous phase, and the percentage (%) variations of the volumes between the different scanning phases were estimated and compared between the two operators. We considered the following conditions: *in vivo* vs *post mortem* (defined as (*post mortem – in vivo)/in vivo*\*100), the *post mortem* vs *ex vivo* – 1month volumes (expressed as (*ex_vivo_1* – *post mortem)/post mortem* *100), the *in vivo* vs *ex vivo* 1 month volumes (expressed as (*ex_vivo_1* – *in vivo)/in vivo* *100) and, finally, *ex vivo* 1 month vs *ex vivo* 12 months (expressed as ((*ex_vivo_12* – *ex_vivo_1*)/ ex_vivo_1*100).

ITK SNAP software (version 3.8.0, 2019) was used as a tool to collect the segmentation and statistical analysis was performed with MATLAB; the statistical significance was defined as an alpha level of alpha = 0.05.

## Results

### Brain segmentation after 12 months in formalin

The segmentations obtained from two operators after 12 months in formalin are reported in Table 1.

**Table 1.** Summary table for the 11 dogs.

|  |  | **O1** |  |  | **O2** |  |  |  |
| --- | --- | --- | --- | --- | --- | --- | --- | --- |
|  |  | *Mean* | *ST* | *CV%* | *Mean* | *ST* |  | *p value* |
|  |  | *(mm<sup>3</sup>)* | *DEV* |  | *(mm<sup>3</sup>)* | *DEV* | *CV%* |  |
| **Ventricles** | **T1** | 1059.27 | 240.87 | 0.22 | 674.9 | 197.52 | 0.25 | **<0.05** |
| **Caudate** | **T2 L** | 576.09 | 56.82 | 0.09 | 600.36 | 71.69 | 0.11 | 0.38 |
| **Caudate** | **T2 R** | 576.77 | 58.46 | 0.10 | 596.72 | 64.26 | 0.10 | 0.31 |
| **Hippocampus** | **T2 L** | 588.06 | 54.16 | 0.09 | 635.3 | 62.15 | 0.09 | 0.07 |
| **Hippocampus** | **T2 R** | 588.61 | 35.57 | 0.06 | 620.04 | 48.87 | 0.07 | 0.10 |
| **Sub Nigra** | **T2 L** | 48 | 11.04 | 0.22 | 50 | 9.73 | 0.19 | 0.64 |
| **Sub Nigra** | **T2 R** | 54.19 | 9.73 | 0.17 | 57.9 | 9.90 | 0.17 | 0.37 |
| **Lat Geniculate** | **T2 L** | 63.55 | 4.78 | 0.07 | 71.2 | 14.83 | 0.20 | 0.11 |
| **Lat Geniculate** | **T2 R** | 67.26 | 6.92 | 0.10 | 74.68 | 11.98 | 0.16 | 0.09 |
| **Med Geniculate** | **T2 L** | 78.19 | 12.85 | 0.16 | 88.64 | 18.22 | 0.20 | 0.13 |
| **Med Geniculate** | **T2 R** | 85.50 | 12.45 | 0.14 | 93.24 | 14.27 | 0.15 | 0.19 |
| **Putamen** | **T2 L** | 76..14 | 11.88 | 0.15 | 77.25 | 13.46 | 0.17 | 0.84 |
| **Putamen** | **T2 R** | 69.96 | 9.50 | 0.13 | 71.65 | 13.93 | 0.19 | 0.74 |
| **Globus Pallidus** | **T2 L** | 55.06 | 9.10 | 0.16 | 60.18 | 11.36 | 0.18 | 0.25 |
| **Globus Pallidus** | **T2 R** | 58.02 | 8.96 | 0.15 | 61.79 | 11.32 | 0.18 | 0.39 |

Estimated volumes of the considered structures. The obtained measurements reported for operator 1 and 2 (O1/O2), expressed the volume averaged across the considered subjects, their standard deviation and CV%. Within this sample the only statistically significant difference between the operators is observed for the ventricles (red).

For each operator (O1/O2), we report the mean and standard deviations of the volumes and their percentage variations expressed as CV%. The ventricles expressed the highest variability within the same operator (22% for O1 and 25% for O2), followed by the Substantia Nigra and Geniculate (around 20%) while the remaining structures showed a variability around 15% or less (see Fig 1).

**Fig 1.**
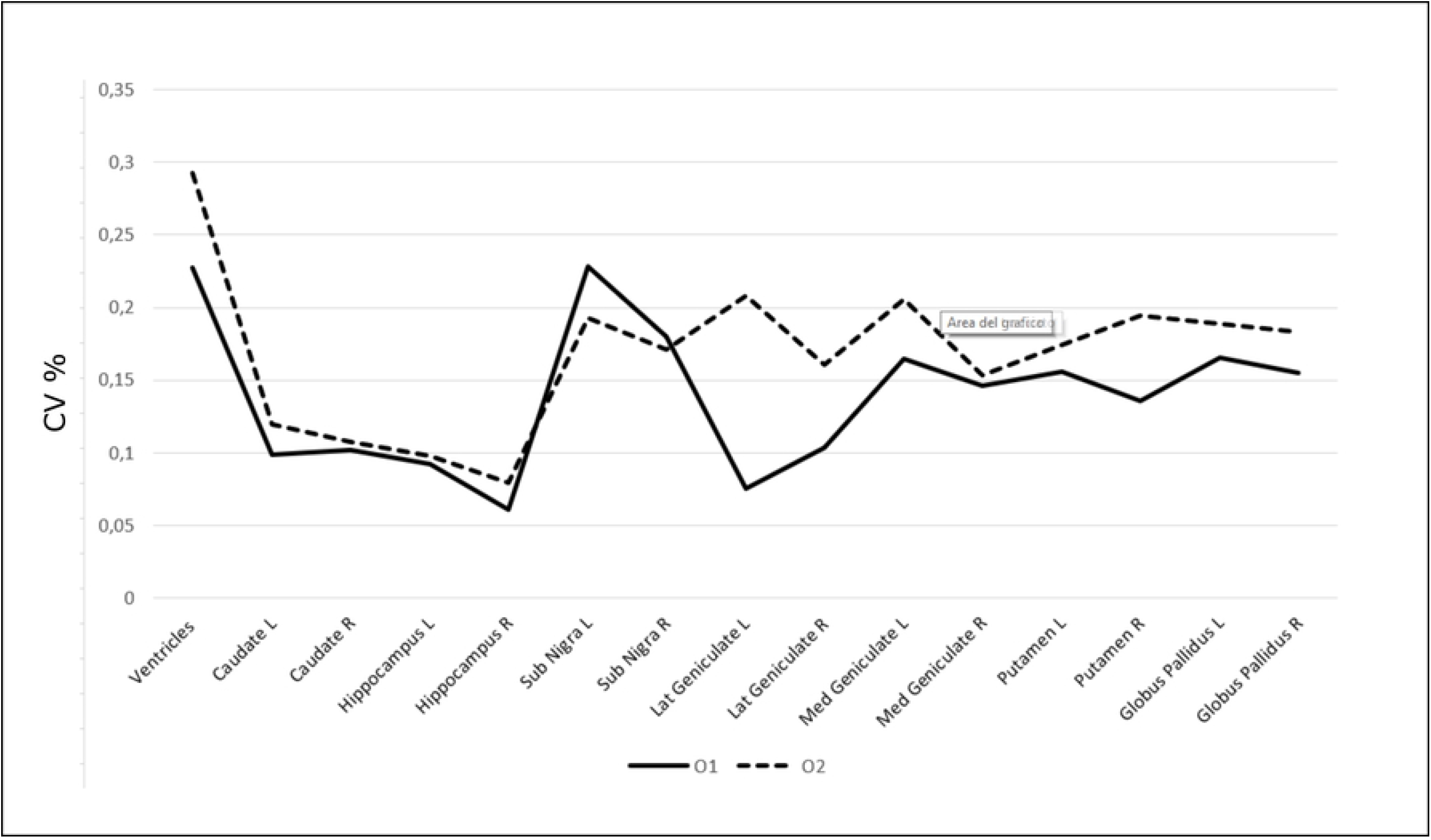
Segmentation stability across operators. The percentage variability of each structure operator O1 (solid line) and O2 (dotted line). It can be noticed that the lowest variability, except for the ventricles, was observed for both operators for the largest and more defined brain structures, thus suggesting that the MRI volume of the segmented structure is influenced by the actual size and its intrinsic contrast with the surrounding parenchyma.

The spatial topography of these variations is shown in Fig 2, where the % CV, averaged across operators, is overlaid on a T1w image of a representative dog.

**Fig 2.**
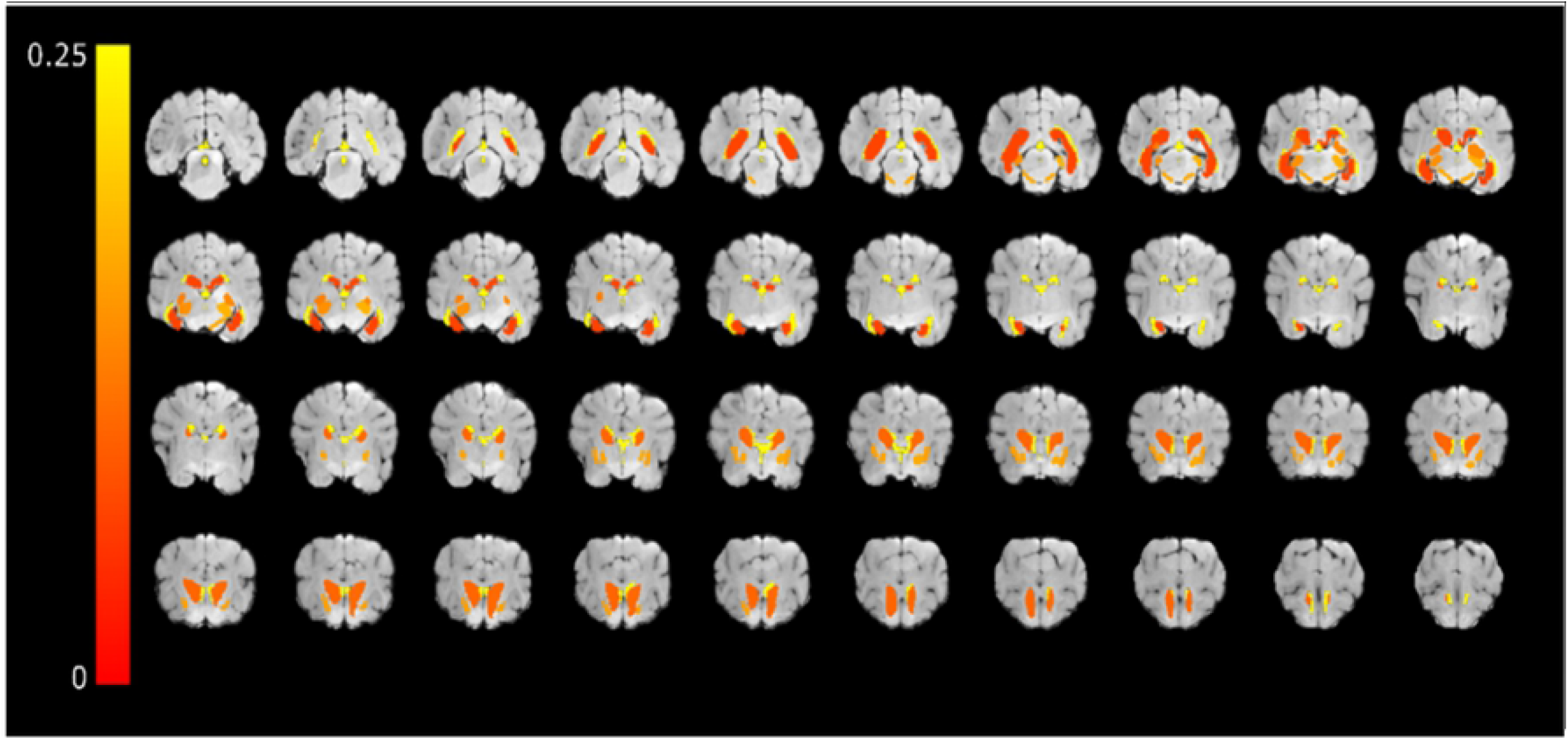
The spatial topography of the % CV of the considered structures. The % CV averaged across operators overlaid on T1w images of a representative dog.

As it can be seen in Table 1 and Fig 3, where the whisker plots of the analyzed structures are reported, apart from the ventricles (p <0.01 – dotted box in Fig 3), a t-test evidenced no statistically significant differences among the two users.

**Fig 3.**
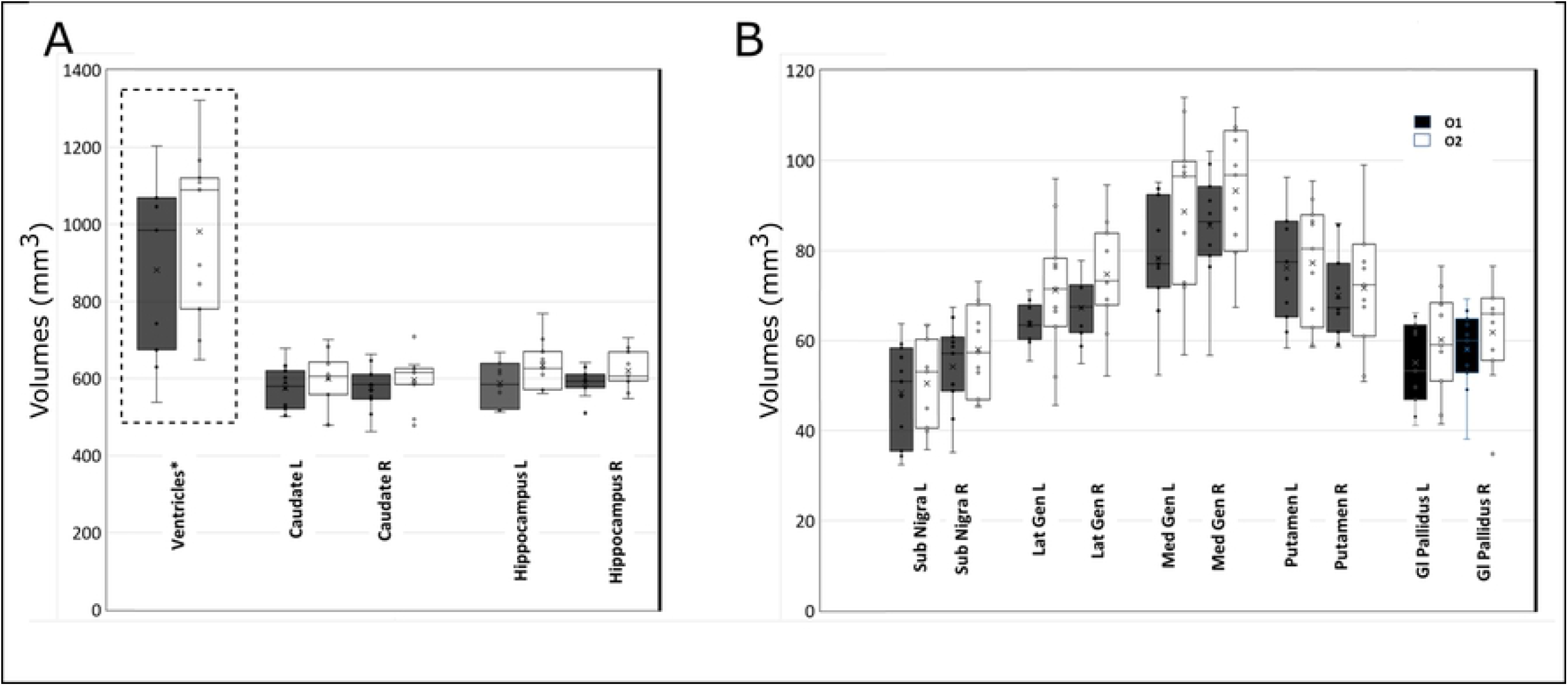
Data distribution of the segmented structures. Whisker plot for the distribution of the segmented volumes from the two operators O1 (black) and O2 (white). A) The set Ventricles, Caudate, and Hippocampus. Only the ventricles resulted statistically different between the two operators (dotted box). B) The set Substantia Nigra, Lateral and Medial Geniculate, Putamen and Globus Pallidus. In this set no statistical difference is observed.

This seems to suggest that the segmentation of the considered structures is quite stable and reproducible across operators. The observed difference in the ventricles might be ascribed to the difficulty to identify their exact border with cerebrospinal fluid (CSF). To summarize, these findings suggest that manual segmentation provides globally reproducible measurements. Further, we observed some structure variability across dogs, that cannot be ascribed to operators’ skills. Such intrinsic structure variability shows that some structures are more stable than others. Of note, larger structures such as Caudate and Hippocampus showed the lowest values of dispersion, while smaller structures such as Medial and Lateral Geniculate were characterized by higher values of dispersion. This might be justified since both operators experienced higher difficulty to segment the smaller structures due to their relatively poor signal contrast.

### A longitudinal study on brain structures

In this part of the study, for a representative dog, we assessed the longitudinal changes of segmentations performed *in vivo, post mortem*, and *ex vivo_1* (after one month in formalin) *and ex vivo_12* (after 12 months in formalin).

First, we observed that both T1w and T2w images showed a change of signal contrast between the *post mortem* and *ex-vivo* data (Fig 4).

**Fig 4.**
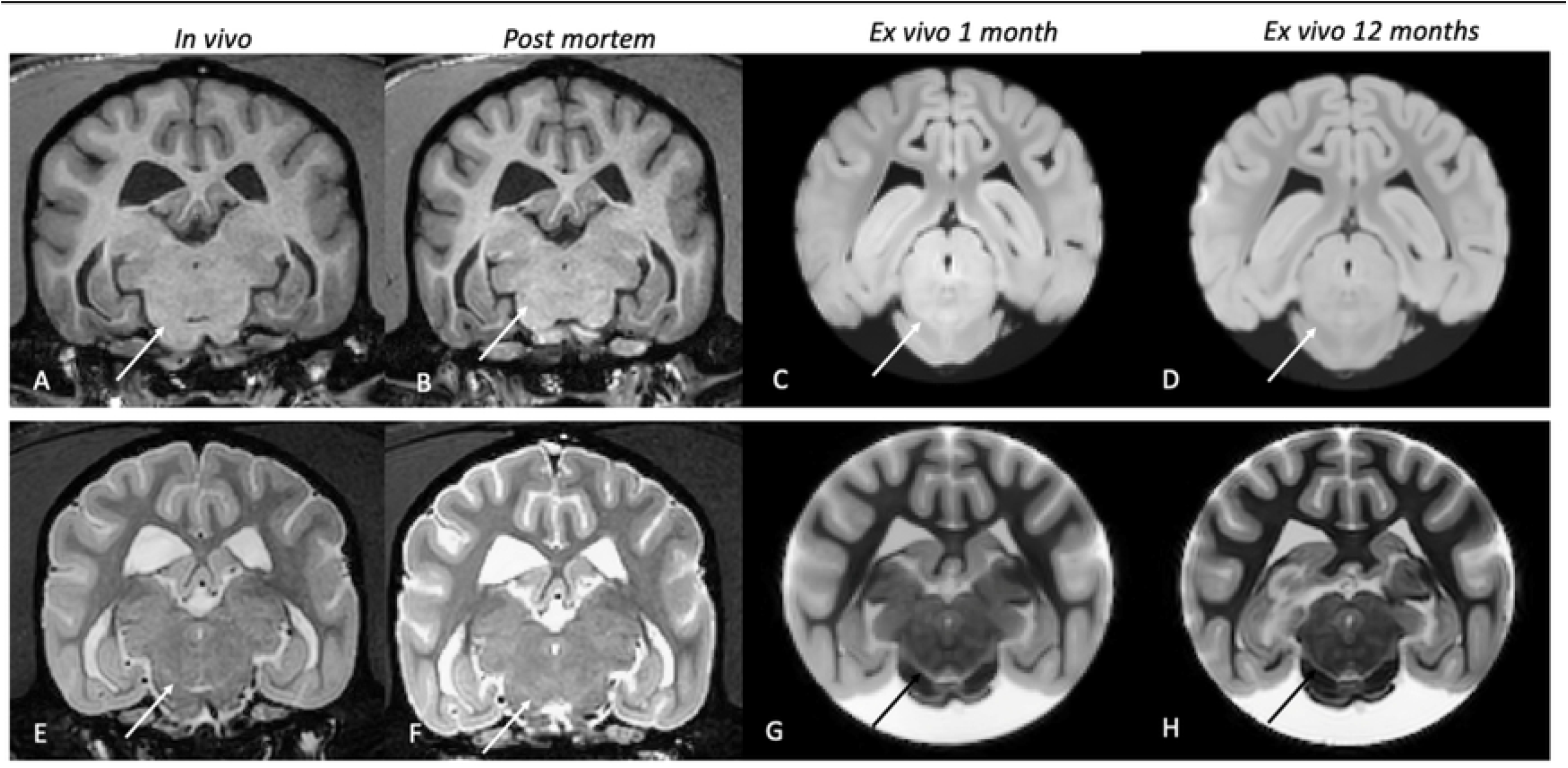
T1 and T2 progressive change of contrast. Panels A-D: Tw transverse section of the studied dog in the four different phases of the experiment, respectively *in vivo* (A), *post mortem* (B), *ex vivo* 1 month (C), and *ex vivo* 12 months (D). *In vivo* and *post mortem* grey and the white matter appeared respectively hypointense and hyperintense, on *ex vivo* images the contrast appeared exactly the opposite, with grey and white matter respectively hyperintense and hypointense. The solid arrows point at the Substantia Nigra and it can be observed the progressive increase of contrast. Panels E-H represent the same four phases from T2w images. In this case, rather than an inversion of the normal contrast, it has been observed a strong decrease of intensity of the white matter in the *ex vivo* phase, thus leading to an increased definition of the contours of the various structures.

Specifically, on T1w images, while *in vivo* (A) and *post mortem* (B) grey and white matter (GM and WM) appeared respectively hypointense and hyperintense, on *ex vivo* images (C-D) the contrast seems the opposite, with GM and WM matter respectively hyperintense and hypointense. This is evident for the Substantia Nigra, (white arrow in the figure). It can be observed that while in panels A and B this structure is barely visible, in panels C (*ex vivo_1*) and D (*ex vivo_12*) the contrast increased with borders of the structure more evident. On T2w images, it is observed a strong decrease in the white matter in both *ex vivo* phases, leading to an increased definition of the contours of the various structures. This is most evident for the smallest structures (e.g. Putamen or Globus Pallidus), whose borders were difficult to define by both operators *in vivo* and *post mortem* phases (Fig 4 E-H). The volumetric percentage variations (see Materials ad Methods) across time of the considered structures are reported in Fig 5.

**Fig 5.**
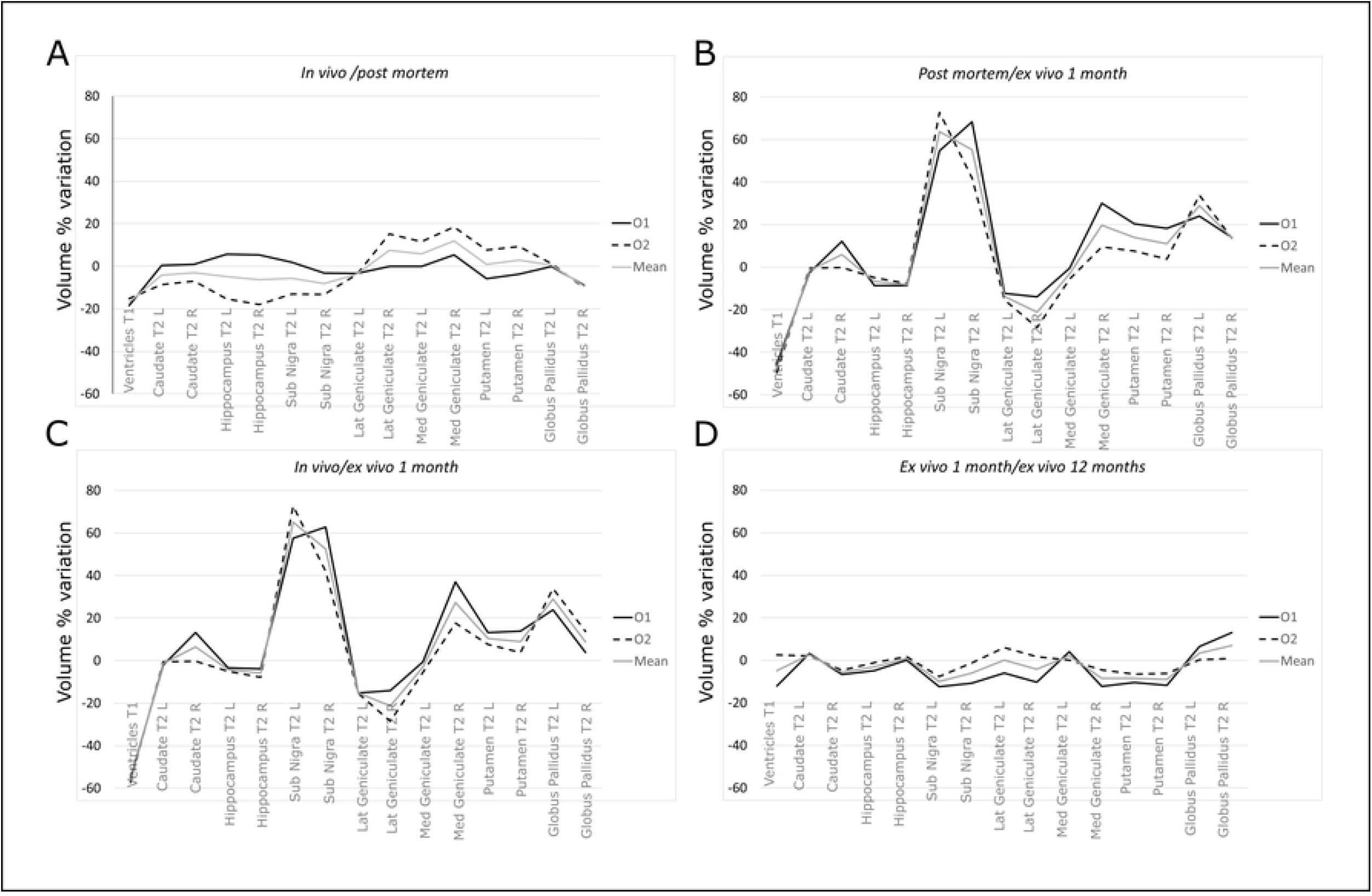
Longitudinal variations. A) Volume variation between the in vivo and post mortem phases. Volume variations were obtained for O1 (solid), O2 (dashed), and the mean trend (gray). It can be observed that the volume variation is always lower than 20%. The trend is variable between the two operators. This aspect may be justified by the difficulty to clearly distinguish the exact borders of the structures in the vivo phase. B) Volume variation between the *post mortem* and *ex_vivo_1* (after 1 month) phases. Volume variations were obtained on each structure for O1 (solid), O2 (dashed), and the mean trend (gray). It can be noted that the agreement between the two operators is higher than in the previous phase, with most structures experiencing an increase in volume from the *post mortem* to *ex vivo* phase. This is particularly interesting since the shift of the contrast after the time in formalin qualitatively seems to improve the visualization of the smallest structures. C) Volume variation between the *in vivo* and *ex_vivo_1* phases. It can be noted that also in this case the operators agree and a trend similar to the previous figure is observed across the structures, i.e. most of the smallest structures appeared more clearly identifiable, thus justifying an increase of the volume. D) Volume variation between the *ex_vivo_1* and *ex_vivo_12* phases. It can be easily observed that the volume variation is relatively low between the two phases, with an overall agreement from both operators, thus suggesting that the time spent in formalin does not significantly influence the volume variation.

As far as it regards the across-operator variability of the segmentations, also in this case, they seem reproducible in the various phases, i.e. the % variation of the volumes globally follows a similar trend for the two operators. Specifically, for the *in vivo/ post mortem* comparison (Fig 5A), the volumetric variation observed in the various structures is close to 12%, fluctuating at most around 20%, and thus can be considered low. Notably, in some structures such as the Lateral and Medial Geniculate nucleus, the volume increased by 18-20%. In this comparison (*in vivo* vs *post mortem*), especially for the *in vivo* images, both operators experienced some difficulties to delineate the borders of the investigated structures. They reported that this was independent of the structures’ dimension, i.e. it applied also to the largest structures. This applies to the in *vivo* phase while in the *post mortem/ ex vivo* comparison, the majority of the structures showed a higher contrast leading to more reproducible segmentations across operators, the highest observed variability ranging around10% (Fig 5B-D). However, we also observed for some structures, a large increase in volume. This was not expected since the formalin fixations are reported to lead to shrinking and reduced tissue volumes (24). This effect was particularly evident for Substantia Nigra (average increase of around 70% Fig 5B, C), followed by Globus Pallidus (around 30%) and Geniculate (around 20%). In general, for both ex vivo phases, all the operators described an increased contrast to detect the borders of the various structures, and the smallest structures were described to be sensitively easier to be segmented. Considering the previous findings, this suggests that the increase of contrast seems to be ascribed to the *ex-vivo* condition and thus the effect of fixation. Now, to study if this change remains stable over time, we compared the two ex vivo conditions (*ex_vivo_1* and *ex_vivo_12*). As it can be seen in Fig 5D, the highest changes fluctuate around 20%. This suggests that the increased time spent in formalin did not influence significantly the volumes of the various structures. Basically, in this period the structures’ volumes remained stable. This is an important finding suggesting that the segmentation can be performed even after a significant amount of time from the formalin fixation. In order to understand if these volume changes were induced by an overall shrinkage or inflation of the brain, we performed the coregistration of the data acquired at the different time points.

As an example, in Fig 6, we report the borders of the brain extracted (for a representative slice) from *in-vivo* data overlaid to the T1w images obtained *ex-vivo* (central panel).

**Fig 6.**
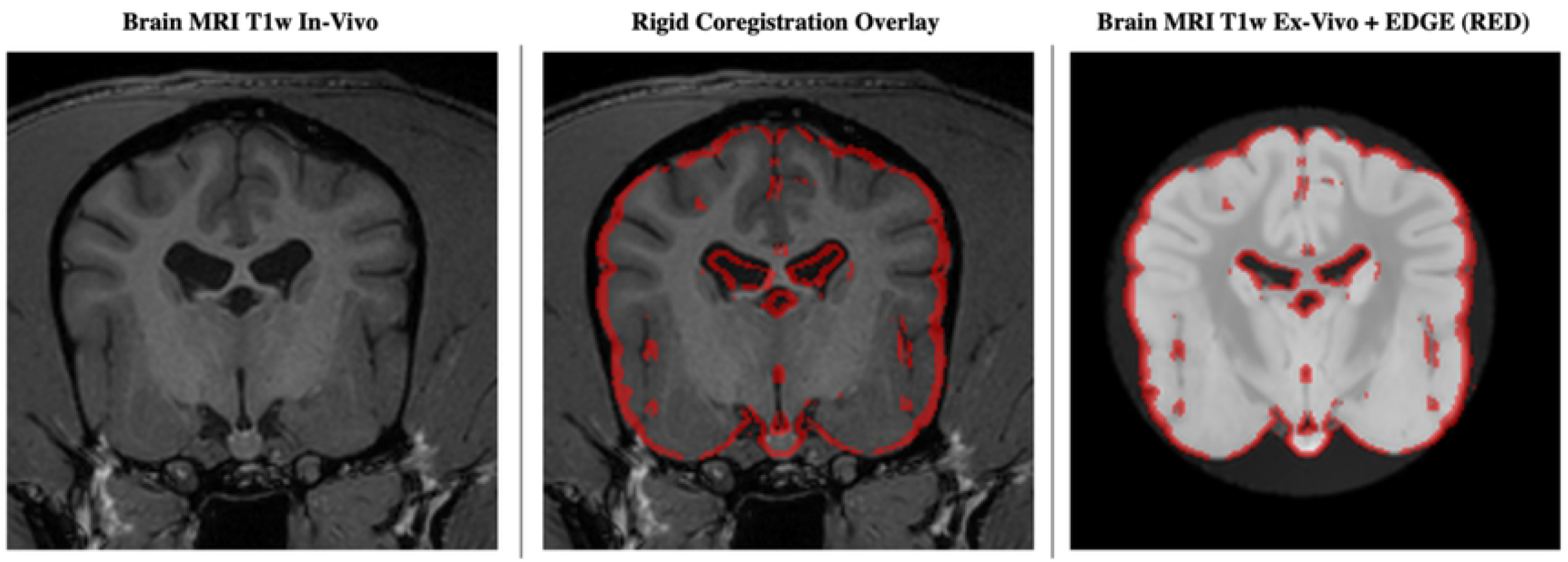
Data coregistration. T1w *in vivo* data are used as the reference to coregister the *ex-vivo* data with a 6-parameter coregistration approach (no scaling included). As an example, a representative slice is reported (A). The *in vivo* brain contours are overlaid to the *ex-vivo* data after the coregistration (B). It can be noted a good agreement. Analogously, the *in-vivo* brain contours overlaid on the *ex vivo* 12 months show that the rigid coregistration successfully aligned the data (C). Since no scaling was involved in the data transformation, this suggests that the brain did not experience any significant inflation/shrinkage.

It can be noted that a rigid co-registration with 6 parameters, thus excluding any scaling factor, was able to coregister the brain successfully. Moreover, as it can be noted in Fig 6 (left panel), the sample applies to the coregistration of the *ex-vivo* data after 12 months. The fact that no scaling factor was needed to coregister the data, seems to suggest that the brain did not experiment with any significant inflation/deflation over time. Therefore, the changes observed in the volumes were likely due to a change in the signal contrast.

So far, the two samples of dogs have been treated separately. However, an interesting point would be if the longitudinal changes observed on a single dog hold also for the sample of 11 dogs. To address this aspect, in a future study we will perform the same longitudinal study on a sample of dogs. However, with the data available at this stage, we tried to assess if there were statistical differences between the volumes obtained from the single dog and the 11-dog sample. To this aim, we considered the distribution of the volumes, structure by structure, obtained from the sample and we tested whether the volumes obtained from the single dog (after 12 months in formalin) belonged to the same distribution or not, i.e. if they were statistically different.

As it can be seen in Table 2, where we report the 95% confidence intervals and the test outcomes, apart from Ventricles, Right Hippocampus and Right Substantia Nigra all structures were not statistically different.

**Table 2.**
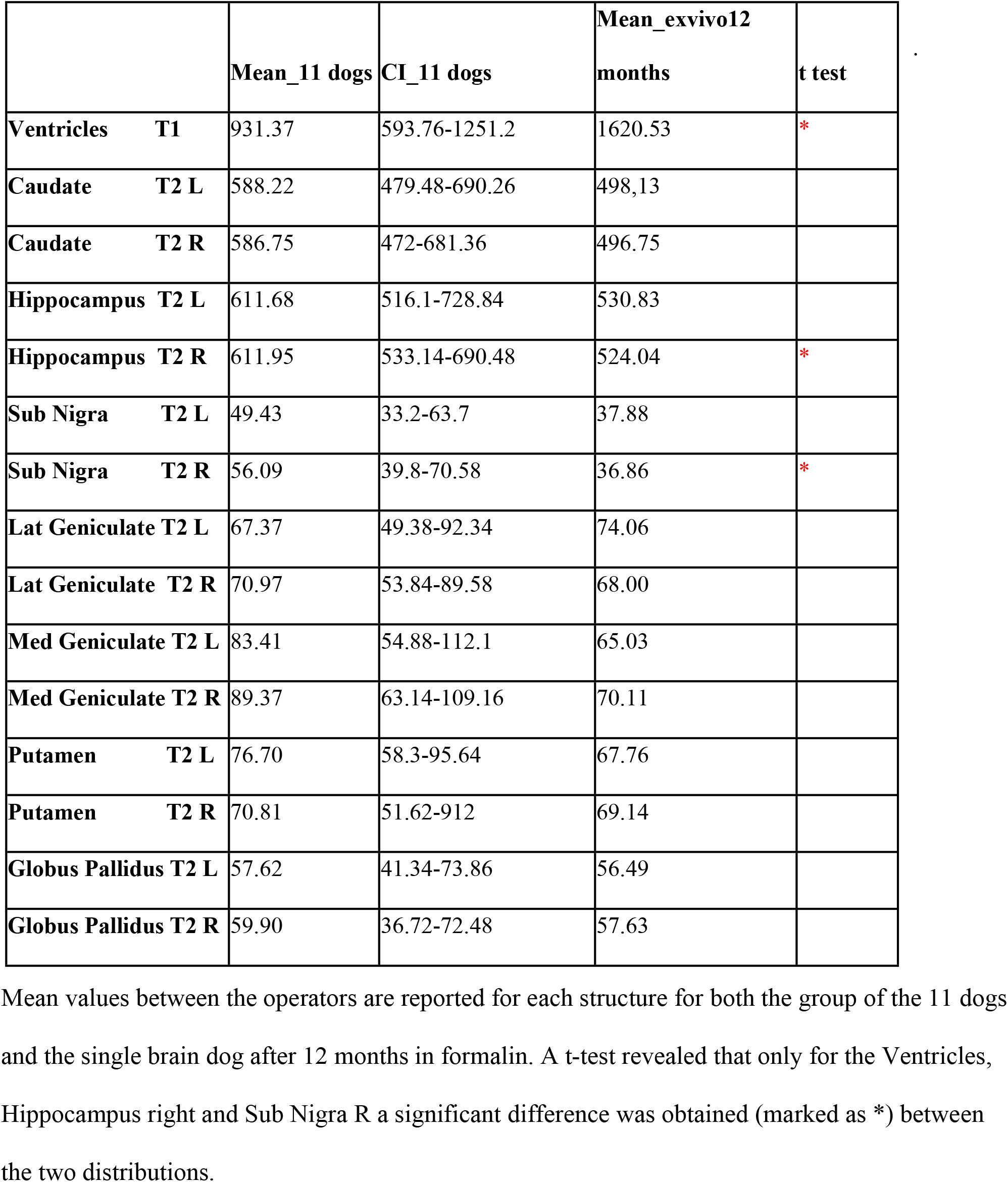
Comparison between the measurements obtained on the group of 11 dogs (entire head) after 12 months in formalin and the ones obtained on the single brain dog after 12 months in formalin.

|  |  | Mean_11 dogs | CI_11 dogs | Mean_exvivo12<br>months | t test |
| --- | --- | --- | --- | --- | --- |
| Ventricles | T1 | 931.37 | 593.76-1251.2 | 1620.53 | * |
| Caudate | T2 L | 588.22 | 479.48-690.26 | 498,13 |  |
| Caudate | T2 R | 586.75 | 472-681.36 | 496.75 |  |
| Hippocampus | T2 L | 611.68 | 516.1-728.84 | 530.83 |  |
| Hippocampus | T2 R | 611.95 | 533.14-690.48 | 524.04 | * |
| Sub Nigra | T2 L | 49.43 | 33.2-63.7 | 37.88 |  |
| Sub Nigra | T2 R | 56.09 | 39.8-70.58 | 36.86 | * |
| Lat Geniculate | T2 L | 67.37 | 49.38-92.34 | 74.06 |  |
| Lat Geniculate | T2 R | 70.97 | 53.84-89.58 | 68.00 |  |
| Med Geniculate | T2 L | 83.41 | 54.88-112.1 | 65.03 |  |
| Med Geniculate | T2 R | 89.37 | 63.14-109.16 | 70.11 |  |
| Putamen | T2 L | 76.70 | 58.3-95.64 | 67.76 |  |
| Putamen | T2 R | 70.81 | 51.62-912 | 69.14 |  |
| Globus Pallidus | T2 L | 57.62 | 41.34-73.86 | 56.49 |  |
| Globus Pallidus | T2 R | 59.90 | 36.72-72.48 | 57.63 |  |

Mean values between the operators are reported for each structure for both the group of the 11 dogs and the single brain dog after 12 months in formalin. A t-test revealed that only for the Ventricles, Hippocampus right and Sub Nigra R a significant difference was obtained (marked as \*) between the two distributions.

The volumes obtained from the single dog, after 12 months in formalin, seem to be consistent with the volumes obtained from the 11 dogs’ sample. The mast majority, namely 80% of the structures, considered for the longitudinal study, belonged to the same distribution of the 11 dogs. This suggests that the considerations on the longitudinal changes observed on a single dog might hold also for the larger sample. Although encouraging, this point needs to be validated with a larger sample in a future study.

## Discussion

This work has been conceived with a dual purpose. First, we assessed the feasibility of brain segmentations on selected structures in the brain kept in formalin for one year, focusing on the reproducibility of the segmentations as a function of the operator and their intrinsic variability within the sample. This study was performed on a homogeneous sample of 11 dogs. Then, one dog (not part of the previous sample) was used for the longitudinal evaluation of the effect of death and fixation on MRI properties of the brain. We obtained that after one year in formalin the segmentations seem reliable and mostly reproducible across operators. Of note, we observed that the time spent in formalin increased the contrast for some specific structures, such as the Substantia Nigra.

The impact of Beagle cerebral models on MRI translational studies is currently increasing. For this reason, several groups are interested in characterizing brain structures. To this aim, in [11,13,14,25] canine MRI based atlases are been developed. However, at the current stage, these are either based on a heterogeneous group of dogs, i.e. a mixed breed sample [3], or on just Beagle dogs, but considering only WM, GM, and CSF, and not specific structures [14]. Further, while these atlases were obtained from alive dogs, in [11] a mixed-breed atlas was obtained by coregistering i*n vivo* with *ex vivo* data. In this case, brains were extracted from the skull and fixed in 10% buffered formalin. Compared to these works, in our study, for the first time to our knowledge, selected brain structures have been segmented on brains fixed in formalin to provide their volumetric evaluation. This allows assessing the reproducibility of such structures, an aspect that has not been previously evaluated. This point is important since manual segmentation is still the more accurate approach, being automatic tools not yet available for these applications [26]. Our findings suggest that the segmentation procedure on this kind of data can be considered overall reproducible since only slight statistical differences were detected, apart from the ventricles. This could be due to the fact that ventricles are fluid filled structures, and they may not be clearly distinguished from the rest of brain CSF.

Furthermore, the fact that some structures are characterized by higher volume variations than others may rely on an intrinsic individual variability. This aspect is particularly interesting, since single structures may be the target of specific experiments, and to determine that there may be an intrinsic variability could be crucial, e.g. in disentangling a pathology/drug induced effect vs a physiological variation. Next to strictly experimental purposes, since the estimation of volumes can assist in the detection and monitoring of the progression of brain disease/damage, the accuracy and reproducibility of MRI segmentation are important also in the clinical field [27]. In fact, nowadays manual segmentation for brain structures or lesions is considered the gold standard in terms of precision, even if it may result in longer analysis times compared to a more “automatic” approach [28,29]. This is in line with [14,22], where the mentioned brain templates were obtained from manually segmented data. Compared to histopathological analyses, the MRI based segmentation of formalin fixed brains has several advantages. First, structural abnormalities can be assessed within the entire brain without altering the original structures. Second, the formalin fixation allows to analyze the data multiple times in any plane to focus on specific areas [19,30] by preserving the structure of the tissues. In fact, after death, the brain undergoes microbial degradation, autolysis, and breakdown of cell membranes. The chemical fixation tends to preserve the macromolecular structure providing the longitudinal stability required for extensive scanning times. Nevertheless, due to death and fixation, a series of artifacts and changes are expected. As far as it regards eventual longitudinal studies, it is known that the formalin fixation may alter the relaxation times. This is due to the induced tissue dehydration, crosslinking, and reduced transmembrane water exchange [30] that lead to a T1/T2 shortening. This results in a higher spatial resolution in terms of borders visualization, for technical details see for example [19,31]. This is in line with what we observed in the *ex vivo* phases, where both operators found an increase of the volumes, especially for the smallest structures, that were judged easier to be segmented. To support this interpretation, we carefully checked the brain volumes by co-registering the brains across the three phases. We obtained no significant changes. Therefore, this apparent increase of volume did not correspond to a real increase in the size of the structure but to an increase of contrast in the structures’ borders allowing a more accurate segmentation. Evidently, before fixation, the structures’ volumes were under-estimated due to low contrast. This is in line with previous findings, see for example [32] where it was reported that the qualitative image evaluation significantly improved after fixation, and the structures segmentations were described to be easier than *in vivo* images.

Another effect of formalin fixation that has been reported in the literature is the tissue shrinkage which may not be homogeneous among the various brain structures, especially when MRI is performed after fixation, see [22]. For example, it has been observed that ventricles may experience filling or emptying according to pressures from the surrounding tissue, and shrinkage of surrounding tissue may not always be paired with an expansion of the ventricles [33] with those tissues experiencing a “positive formalin effect”, characterized by a swelling effect caused by the osmotic pressure of the formalin solution [34]. These aspects should be taken into proper considerations when volumetric evaluations are performed on brains in *post mortem* or *ex vivo* conditions. These effects may relate to the observed variability of the ventricles in our data.

Finally, according to our findings, the amount of time (12 months) spent in formalin seems not to influence the volumes of the structures: their percentage variation did not exceed 10%. This aspect is important since it suggests that the same brain can be potentially used for several studies, even after some time, without the risk of significant structural changes. This observation is consistent with data proposed by other studies since previous investigations on the relationship between MR volumetric measurements performed in-vivo and ex-vivo report that the brain structures remain relatively stable for 6 months post-mortem [35]. This applies also to human medicine where it has been reported that fixation leads to no significant leaching of iron in long term storage [36] and that WM components, including myelin, are all well preserved [37].

To summarize, based on our findings *post mortem* MRI based segmentation seems to be a useful and accurate tool that allows longitudinal studies. However, especially for the observed longitudinal variations, these findings need to be further validated on a larger sample.

